# Isolation of infective Zika virus from urine and saliva of patients in Brazil

**DOI:** 10.1101/045443

**Authors:** Myrna C. Bonaldo, Ieda P. Ribeiro, Noemia S. Lima, Alexandre A. C. dos Santos, Lidiane S. R. Menezes, Stephanie O. D. da Cruz, Iasmim S. da Mello, Nathália D. Furtado, Elaine E. de Moura, Luana Damasceno, Keli A. B. da Silva, Marcia G. da Castro, Alexandra L. Gerber, Luiz G. P. da Almeida, Ricardo Lourenço-de-Oliveira, Ana Tereza R. Vasconcelos, Patrícia Brasil

## Abstract

**BACKGROUND:** Zika virus (ZIKV) is an emergent threat provoking a worldwide explosive outbreak. Since January 2015, 41 countries reported autochthonous cases. In Brazil, an increase in Guillain-Barré syndrome and microcephaly cases was linked to ZIKV infections. A recent report describing low experimental transmission efficiency of its main putative vector, *Ae. aegypti,* in conjunction with apparent sexual transmission notifications prompted the investigation of other potential sources of viral dissemination. Urine and saliva have been previously established as useful tools in ZIKV diagnosis. However, no evidence regarding the infectivity of ZIKV particles present in saliva and urine has been obtained yet.

**METHODOLOGY/PRINCIPAL FINDINGS:** Nine urine and five saliva samples from nine patients from Rio de Janeiro presenting rash and other typical Zika acute phase symptoms were inoculated in Vero cell culture and submitted to specific ZIKV RNA detection and quantification through, respectively, NAT-Zika, RT-PCR and RT-qPCR. Two ZIKV isolates were achieved, one from urine and one from saliva specimens. ZIKV nucleic acid was identified by all methods in four patients. Whenever both urine and saliva samples were available from the same patient, urine viral loads were higher, corroborating the general sense that it is a better source for ZIKV molecular diagnostic. In spite of this, from the two isolated strains, each from one patient, only one derived from urine, suggesting that other factors, like the acidic nature of this fluid, might interfere with virion infectivity. The complete genome of both ZIKV isolates was obtained. Phylogenetic analysis revealed similarity with strains previously isolated during the South America outbreak.

**CONCLUSIONS/SIGNIFICANCE:** The detection of infectious ZIKV particles in urine and saliva of patients during the acute phase may represent a critical factor in the spread of virus. The epidemiological relevance of this finding, regarding the contribution of alternative non vectorial ZIKV transmission routes, needs further investigation.

**AUTHOR SUMMARY:** The American continent has recently been the scene of a devastating epidemic of Zika virus and its severe manifestations, such as microcephaly in newborns and Guillain-Barré Syndrome. Zika virus, first detected in 1947 in Africa, only from 2007 started provoking outbreaks. Zika, dengue and chikungunya viruses are primarily transmitted by *Aedes* mosquitoes. Dengue is endemic in Brazil for almost 30 years, and the country is largely infested by its main vector, *Aedes aegypti*. Chikungunya virus entered the country in late 2014 and Zika presence was confirmed eight months later. Nevertheless, Zika notifications multiplied and spread across the country with unprecedented speed, raising the possibility of other transmission routes. This hypothesis was strengthened by some recent reports of Zika sexual transmission in *Ae. aegypti*-free areas and by the description of a low transmission efficiency to Zika virus in local *Ae. aegypti*. We found Zika active particles in both urine and saliva of acute phase patients, and a finding that was promptly announced by Fiocruz via Press Conference on February 5, 2016. In this work, we bring up the potential alternative person-to-person infection routes beyond the vectorial transmission, that might have epidemiological relevance.

## INTRODUCTION

Zika virus (ZIKV) is an emerging mosquito-borne virus of the family *Flaviviridae* and genus *Flavivirus* [1]. ZIKV was first reported in 1947 after isolation from a febrile sentinel rhesus monkey [2]. Since then, serologic evidence of human ZIKV infection in Africa and Asia was detected, but until 2005 only few human cases were reported [3]. The first well-described outbreak outside these geographic regions happened in 2007 in Micronesia, more specifically in Yap State, when the majority of the population was affected with Zika fever [4]. Intriguingly, the local mosquito vector was not confirmed by neither viral isolation nor molecular methods [4].

On October 2013, a second intense outbreak in Oceania occurred in French Polynesia (2013/2014), and soon after spread over to New Caledonia (2014), Cook Islands, (2014) and Ester Island, 2014 [5, 6]. In these outbreaks, approximately 80 % of ZIKV infections was asymptomatic [4, 7]. Commonly, Zika is considered to be a mild disease lasting one week with symptoms including fever, rash, conjunctivitis, arthralgia, myalgia, headache and malaise. However, during the French Polynesian epidemic, its association with severe neurological complications, the Guillain-Barré syndrome (GBS) was reported for the first time [8].

In April 2015, the first autochthone cases in the Americas was identified in Brazil [9, 10]. At present, Brazil suffers an explosive outbreak of ZIKV. Hence, in February 2016, Brazilian Ministry of Health (MoH) appraised the incidence of further than one million of ZIKV disease cases [11]. Notably, in addition of an increase of GBS cases as occurred in French Polynesia outbreak, the MoH of Brazil described a rise of microcephaly occurrence. Between 22 October 2015 to 5 March 2016, 6158 cases of microcephaly and/or central nervous system malformation were noticed in contrast to the estimated average number of 163 annual cases [12]. So far, 745 suspected cases of microcephaly have been confirmed as ZIKV-associated microcephaly in a total of 1927 investigated cases [11-13]. More recently, a case of ZIKV infection with vertical transmission demonstrated the association of severe fetal brain injury with fetal infection with ZIKV [14]. Moreover, ZIKV nucleic acid was detected in amniotic fluid of two pregnant women, whose fetuses were diagnosed with microcephaly, corroborating vertical transmission possibility [15]. Other abnormalities such as placental insufficiency, fetal growth restriction, CNS injury, and fetal death have also been reported in association with ZIKV infection [16]. This scenario of ZIKV infection linked to severe neurological complications as well as the establishment of ongoing ZIKV outbreaks in several countries in Latin America led to the WHO to declare ZIKV an international public health emergency [11, 17, 18].

The transmission of ZIKV has been associated with several *Aedes* mosquito species belonging to subgenus *Stegomyia*, notably *Ae. aegypti* [19, 20] and *Ae. albopictus* [21]. However, a recent study proposes that although susceptible to infection, *Ae. aegypti* and *Ae. albopictus* from the Americas display an unexpectedly low vector competence for ZIKV [22], suggesting other factors such as the large naïve population for ZIKV and the high densities of human-biting mosquitoes contribute to the rapid spread of ZIKV during the current outbreak. Nonetheless, perinatal transmission [23] and potential risk for transfusion-transmitted ZIKV infections has also been demonstrated [24]. Most remarkably, ZIKV can be likely disseminated by sexual contact, due to its presence in semen [25, 26]. In addition, it was demonstrated the existence of ZIKV in urine [27, 28], breast milk [29] and saliva [30]. Indeed, ZIKV was more frequently detected in urine and saliva than in blood using ZIKV RT-PCR tests for diagnosis. It was considered that patients exhibit the highest concentrations of ZIKV in saliva at disease onset [30] while in urine, ZIKV possibly remains detectable for longer periods [27]. So far, no evidence has been obtained regarding the infectivity of ZIKV particles present in saliva and urine. In this study, we demonstrate that it is possible to recover infective ZIKV from both saliva and urine of acute phase patients by means of viral isolation in Vero cells. This achievement suggest that ZIKV may be transmitted between humans by infected saliva and urine.

## METHODS

### Study facilities and patients enrollment

The Acute Febrile Illnesses Laboratory and Molecular Biology of Flavivirus Laboratory conducted this study at Oswaldo Cruz Foundation, Rio de Janeiro. The institutional review boards at Fundação Oswaldo Cruz (Fiocruz) approved the study protocol. All subjects provided written, informed consent before participation, and a medical assistant filled a standardized medical questionnaire form, during an interview with the participants. Numbers of days from the first reported symptom (days after symptoms onset) and main signs and symptoms were recorded. Urine and saliva samples investigated in this study were collected from January 14^th^ to February 2^nd^, 2016. The samples were obtained only from patients clinically suspected with ZIKV infection, especially with pruritic maculopapular rash.

### Clinical samples

Saliva and urine specimens were collected in 50 mL sterile certified, DNase-/RNase-free tubes, and after collection, in some cases, the pH was measured by a digital pH meter, in order to investigate the relevance of the pH for viral infection. Twenty five millimeter diameter sterile syringe filters with a 0.22 μm pore size were used to filter the specimens. The samples were aliquoted for subsequently analysis and assays, as infection in Vero cell culture and RNA isolation.

### Primary viral isolation

The African green monkey kidney (Vero) cell line obtained from (ATCC) was grown in 37° C, under an atmosphere containing 5% CO_2_, in Earle’s 199 medium supplemented with 5% fetal bovine serum (FBS). The Vero cells were seeded at a density of 40,000 cells/cm^2^ in 25 cm^2^ culture flasks 24 hours before inoculation. The urine and saliva samples were diluted in Earle’s 199 medium supplemented with 5% FBS (1:2 and 1:4), and 1 mL of each dilution was inoculated onto Vero cells monolayer. After 1 h incubation at 37°C, the inoculum was removed and replaced by 10 mL culture medium. As negative control for each experiment, Vero cells seeded in one culture flask were mock inoculated with culture media. The presence of infectious viral particles was controlled by observation of cytopathic effects (CPE).

### Plaque forming unit assay

Vero cells were seeded at a density of 40,000 cells/cm^2^ in 6-well plates 24 h before inoculation. Dilutions of the biological specimens (1:2, 1:4 and 1:8) in culture media were used to infect monolayers (200 μL/well). After 1 h incubation at 37° C, the inoculum was removed and replaced by 3 mL of 2.4 % CMC (carboxymethyl cellulose) in Earle’s 199 medium. After 7 days incubation at 37° C, cells were fixed with 10 % formaldehyde, washed, and stained with 0.4 % crystal violet for visualization of plaques.

### RNA isolation

Viral RNA was isolated from 140 μL of each biological specimens and cell culture supernatant using the QIAamp Viral RNA Mini Kit (Qiagen, Hilden, Germany) according to the manufacturer’s recommendations. RNA was eluted in 60 μl of AVE buffer and stored at −80°C until use. The concentration and purity of each RNA sample were measured by Thermo Scientific NanoDrop 8000 Spectrophotometer and Agilent 2100 Bioanalyzer using the Agilent RNA 6000 Nano Kit according the manufacturer’s instructions.

### RT-PCR

The viral RNA was reverse transcribed applying the Superscript IV First-Strand Synthesis System (Invitrogen) using random hexamers according to the manufacturer’s recommendations. The reverse transcription reaction was carried out at 23°C for 10 min, 55°C for 10 min and 80°C for 10 min. Further, the viral RNA was amplified by conventional PCR covering viral NS1 genomic region (genome position 3085-3385) using GoTaq Green Master Mix (Promega) according to the manufacturer’s recommendations. The thermocycling program set up in a Veriti 96 Well thermocycler (Applied Biosystem) was 1 cycle of 95°C for 5min; 40 cycles of 95°C for 40 sec, 50°C for 40 sec, 72°C for 30 sec; 1 cycle of 72°C for 10min and hold of 4°C. 10 ml of Amplified products were detected by electrophoresis on a 2% agarose gel, visualized by ethidium bromide staining UV.

### NAT and Quantitative RT-PCR

To discard co-infection of ZIKV with dengue and/or chikungunya viruses, we analyzed the urine, saliva samples and the viral strains isolated from Vero cell using he NAT- Dengue, Zika and Chikungunya discriminatory kit (Instituto de Biologia Molecular do Paraná and Fundação Oswaldo Cruz, Brazil). To measure genomic ZIKV load, viral RNA was reverse transcribed and amplified using the TaqMan Fast Virus 1-Step Master Mix (Applied Biosystems) in an Applied Biosystems StepOnePlus Instrument. For each reaction we used 400 nM forward primer (5’-CTTGGAGTGCTTGTGATT-3’, genome position 3451-3468), 600 nM reverse primer (5’-CTCCTCCAGTGTTCATTT-3’, genome position 3637-3620) and 250 nM probe (5’FAM-AGAAGAGAATGACCACAAAGATCA-3’TAMRA, genome position 3494-3517). The sequences of this primer set were kindly provided by Isabelle Lepark-Goffart (French National Reference Centre for Arboviruses, IRBA, Marseille, France). Samples were run in duplicate. The reverse transcription was performed at 50°C for 5 minutes. The qPCR conditions were 95°C for 20 seconds, followed by 40 amplification cycles of 95°C for 15 seconds and 60°C for 1 minute. Copy numbers of ZIKV genomic RNA were calculated by absolute quantitation using a standard curve for each run. To construct a standard curve, we cloned an amplicon comprising the genomic region 3085-4032 of the isolate Rio-U1 using pGEM-T Easy Vector (Promega) to serve as a template for in vitro transcription. The RNA transcript was made with mMessage mMachine High Yield Capped RNA Transcription Kit (Invitrogen) using T7 enzyme and purified using MEGAclear Kit (Ambion) according to manufacturer’s instructions. The purity of the transcript was verified using NanoDrop 8000 Spectrophotometer (Thermo Scientific), the integrity was analyzed using 2100 Bioanalyzer (Agilent) using the RNA 6000 Nano Kit (Agilent), and the concentration of the RNA was accessed using Qubit 2.0 Fluorometer (Invitrogen). The standard curve was generated by a ten-fold dilution (ranging from 10 to 10^9^ copies/reaction) of the transcript. The limit of detection under standard assay conditions was approximately 40 viral RNA copies/mL.

### Nucleotide Sequence

Double-stranded cDNA libraries were prepared using the TruSeq Stranded mRNA LT Sample Preparation Kit (Illumina, San Diego, CA, USA). Briefly, the polyA containing mRNA purification step was not performed and the protocol was started with 25-35 ng of RNA in 5 ul of molecular biology grade water to which were added 13 ul of Fragment, Prime, Finish Mix. The remaining steps of the protocol were carried out without any modifications. Library quality control was performed using the 2100 Bioanalyzer System with the Agilent DNA 1000 Kit (Agilent, Santa Clara, CA, USA). The libraries were individually quantified via qPCR using a KAPA Library Quantification Kits for Illumina platforms (KAPA Biosystems, Wilmington, MA, USA). The libraries were pooled together in equimolar quantities and sequenced. Paired-end reads (2 × 75 bp) were obtained using a MiSeq Reagent Kits v3 (150-cycles) in a MiSeq sequencing system (Illumina).

### Assembly and annotation

A total of 17,413,830 reads was generated for Rio-U1 sample and 21,734,486 for Rio-S1 sample. Related reads to *Chlorocebus sabaeus* have been filtered using Bowtie2 and Samtools, remaining 12,614,062 reads of Rio-U1 and 12,943,134 of Rio-S1. Both genomes were assembled using Ray 2.20 (k = 31). The completed genome of Rio-U1 has 10,795bp (Acession number KU926309) and Rio-S1 has 10,805bp (Acession number KU926310). Gene prediction was performed by GenemarkS 4.17. Mature peptides were identified by blastp against the protein annotated in reference sequence NC_012532.

### Phylogenetic analysis

Phylogenetic reconstruction was inferred by means Maximum Likelihood method based on the General Time Reversible model with a discrete Gamma distribution and invariable sites (GTR+G+I) using alignments of poliprotein genes from 39 Zika virus strains and 1 dengue virus serotype 4.

## RESULTS

### Patients, clinical and social demographic characteristics

We examined nine enrolled patients suspected of ZIKV infection. The initial medical support and collection of urine and saliva samples were performed from January 14^th^ to February 2^nd^ 2016. Out of seven women, six were pregnant with gestational ages varying from 18 to 33 weeks, median value of 20. 5 ± 5.8 weeks (Supplementary Table 1 and 2). The female patient ages ranged from 20 to 42 years old (median value of 28.5 ±7.4 years) and the male patient ages were 24 and 68 years old. All the patients live in the metropolitan area of Rio de Janeiro (Supplementary Tables 3 and 4).

**Table 1.**
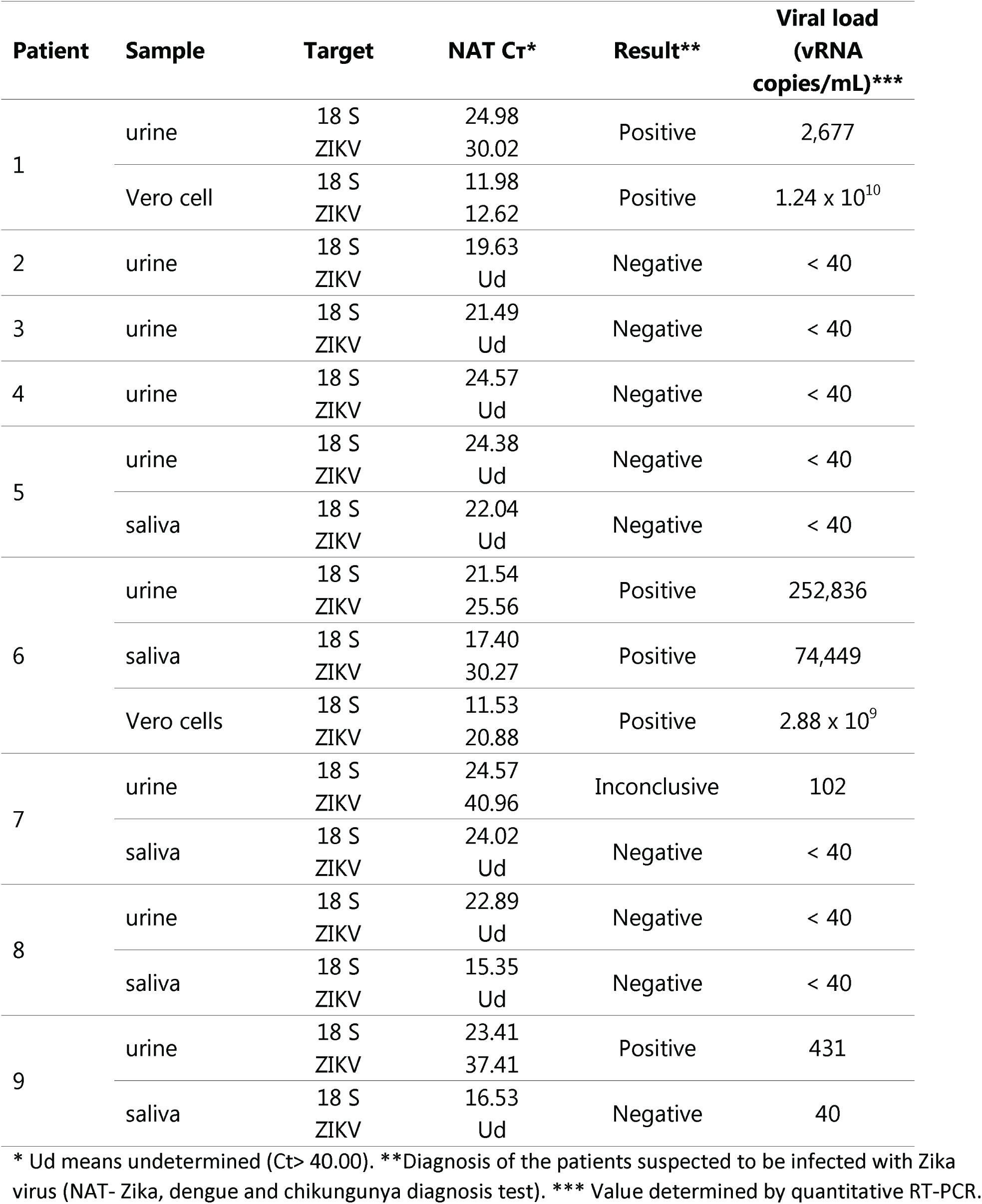
ZIKV RNA detection and quantitation

| Patient | Sample | Target | NAT Ct* | Result** | Viral load<br>(vRNA<br>copies/mL)*** |
| --- | --- | --- | --- | --- | --- |
| 1 | urine | 18 S | 24.98 | Positive | 2,677 |
|  |  | ZIKV | 30.02 |  |  |
|  | Vero cell | 18 S | 11.98 | Positive | 1.24 x 10<sup>10</sup> |
|  |  | ZIKV | 12.62 |  |  |
| 2 | urine | 18 S | 19.63 | Negative | < 40 |
|  |  | ZIKV | Ud |  |  |
| 3 | urine | 18 S | 21.49 | Negative | < 40 |
|  |  | ZIKV | Ud |  |  |
| 4 | urine | 18 S | 24.57 | Negative | < 40 |
|  |  | ZIKV | Ud |  |  |
| 5 | urine | 18 S | 24.38 | Negative | < 40 |
|  |  | ZIKV | Ud |  |  |
|  | saliva | 18 S | 22.04 | Negative | < 40 |
|  |  | ZIKV | Ud |  |  |
| 6 | urine | 18 S | 21.54 | Positive | 252,836 |
|  |  | ZIKV | 25.56 |  |  |
|  | saliva | 18 S | 17.40 | Positive | 74,449 |
|  |  | ZIKV | 30.27 |  |  |
|  | Vero cells | 18 S | 11.53 | Positive | 2.88 x 10<sup>9</sup> |
|  |  | ZIKV | 20.88 |  |  |
| 7 | urine | 18 S | 24.57 | Inconclusive | 102 |
|  |  | ZIKV | 40.96 |  |  |
|  | saliva | 18 S | 24.02 | Negative | < 40 |
|  |  | ZIKV | Ud |  |  |
| 8 | urine | 18 S | 22.89 | Negative | < 40 |
|  |  | ZIKV | Ud |  |  |
|  | saliva | 18 S | 15.35 | Negative | < 40 |
|  |  | ZIKV | Ud |  |  |
| 9 | urine | 18 S | 23.41 | Positive | 431 |
|  |  | ZIKV | 37.41 |  |  |
|  | saliva | 18 S | 16.53 | Negative | 40 |
|  |  | ZIKV | Ud |  |  |

\* Ud means undetermined (Ct> 40.00). \*\*Diagnosis of the patients suspected to be infected with Zika virus (NAT- Zika, dengue and chikungunya diagnosis test). \*\*\* Value determined by quantitative RT-PCR.

**Table 2.** Differences in amino acid residues in ZIKV polyproteins of Rio-Sl and Rio-U1 isolates

**Differences in amino acid residues in ZIKV polyproteins of Rio-S1 and Rio-U1 isolates**
| Polyprotein position | ZIKV protein (amino acid position) | Rio-S1 | Rio-U1 |
| --- | --- | --- | --- |
| 625 | E (335) | A | T |
| 1143 | NS1 (349) | V | M |
| 1404 | NS2B (32) | M | I |
| 2039 | NS3 (537) | K | R |
| 2122 | NS4A (3) | T | A |
| 2688 | NS5 (168) | A | V |

The most frequent sign of ZIKV disease was pruritic maculo papular rash which lasted in average 4 days (Supplementary Tables 1 and 2). However, other clinical symptoms were also prevalent, such as low grade fever (< 38°C), headache, myalgia and arthralgia of large and small joints, present in 5 out of 9 patients.

We collected and analyzed urine from patients 1 to 4 and both urine and saliva samples from patients 5 to 9. Vero cells cultures were inoculated at the same date of sample collection and then daily observed through inverted microscopic examination until the appearance of cytopathic effect (CPE). Within one week of incubation, only two samples exhibited CPE (2 out of 14), the urine sample of patient 4 with CPE detected at 4^th^ day of post-inoculation (1 out of 9) and the saliva sample of patient 6 at 5^th^ day post-inoculation (1 out of 5). In this last infection, we recognized small foci of rounded and refractive cells detaching from the monolayer (Fig 1 A and B). After one-week incubation, we proceeded to split cells from negative cultures by means of trypsinization when monolayer was confluent. This procedure was repeated for three consecutive times. Nevertheless, it was not possible to isolate ZIKV in these samples, neither by detecting CPE in Vero cell monolayers or ZIKV genome by RT-PCR (results not shown). We also analyzed these samples by plaque forming assay as a way to detect infectious virus particles. Unfortunately, we did not perform this analysis with urine of patient 1, because we received a small aliquot of this specimen. Nevertheless, we detected viral plaques from samples of patient 6 (Fig 1 C and D), in which the dilution 1:2 of saliva originated in 8 PFU resulting in an original viral concentration of 80 PFU/ml in saliva of patient 6. Interestingly, only one viral plaque was visualized by means of this methodology in urine sample of this patient 6, resulting in a titer of 10 PFU/ml.

**Fig 1.**
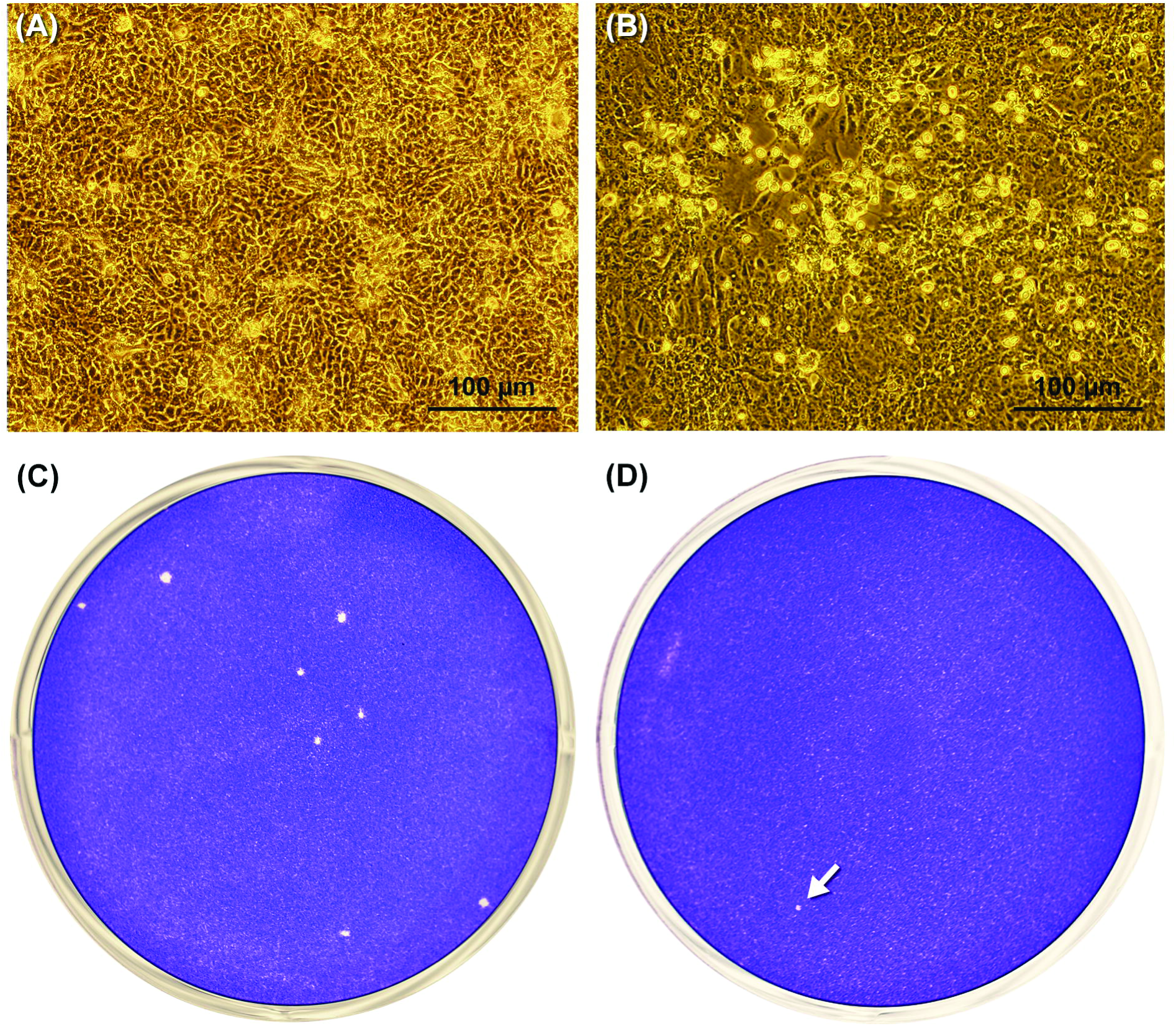
Isolation of Zika virus in Vero cell from the saliva of patient 6.

Phase contrast optical microscopy of culture flasks containing (A) Mock-infected Vero cells and (B) saliva-infected Vero cells presenting a clear visible cytopathic effect. Viral plaque detection in saliva (C) and urine (D). The white arrow shows the unique viral plaque detected in the urine sample.

### ZIKV diagnosis and genome detection

Furthermore, we analyzed all urine and saliva specimens by RT-PCR to confirm the detection of ZIKV (Fig 2A and B). In addition, we included RNA samples of ZIKV isolated from patient 1 and 6 in Vero cells. The set of samples of patient 1 and 6 were all positive and an expected-amplicon band of around 300 bp was seen in electrophoretic analyses, demonstrating the presence of ZIKV genome in these samples (Fig 1 A and B). We also observed a fade band from urine and saliva of patient 9 (Fig 2 B). The ZIKV specificity of this approach was confirmed when we tested this protocol in RNA samples of Chikungunya (CHIKV), dengue (DENV) and yellow fever (YFV) viruses (Fig 2 C).

**Fig 2.**
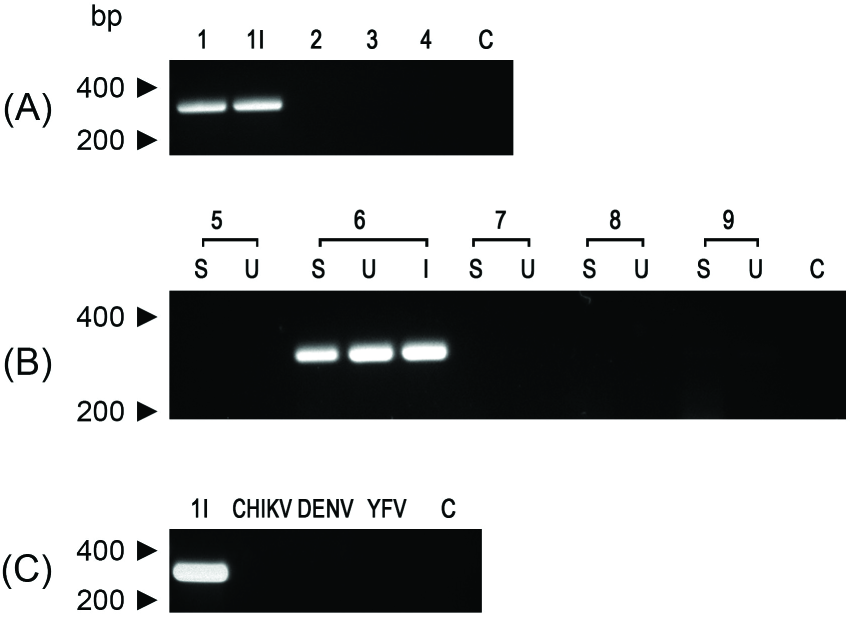
Detection of genomic RNA of Zika virus in urine and saliva samples by RT-PCR analysis. (A) Shows the profiles obtained from urine samples. The lane numbers indicate the patient code. The lane 1I is the amplicon obtained from the viral isolate from urine of patient 1 (isolate Rio-U1). (B) RT-PCR analysis from patients from 5 to 9 where S indicates saliva RNA samples and U, urine RNA samples. (C) Amplification of Zika virus genome of isolate Rio-U1 (1i) with ZK-specific primers that were too employed in the RT-PCR assay of Chikungunya virus RNA (CHIKV), dengue virus RNA (DENV) and Yellow Fever 17DD RNA (YFV). In all of these analyses, a negative control of amplification were included (C). The size marker migration is indicated on the left of the figures.

Notwithstanding, it was mandatory to confirm the result of Zika virus infections in patients and isolations in Vero cells, since ZIKV, DENV and CHIKV are co-circulating in Brazil and the diseases caused by them exhibit similar symptoms. So, each sample was tested for the presence of these three viruses by the ZIKV nucleic acid testing (NAT) of samples which was established to be routinely used in Brazil as diagnosis test since December 2015. (Table 1). All patients included in this study were negative for DENV and CHIKV (Ct > 40.0). Patient 1 was positive for ZIKV in urine (Ct of 30.02) and patient 6 in urine (Ct of 25.56) and saliva (Ct of 30.27) and the viral isolates derived obtained from specimens of these patients were also positive and presented Ct of 12.62 and Ct of 20.88, respectively. Patient 9 was also positive for ZIKV in urine specimen (Table 1), whereas urine from patient 7 presented amplification in a late cycle and, therefore, this result was considered inconclusive. To validate negative results, the ribosomal 18S RNA was detected in all samples showing that there was no inhibition of the RT-PCR.

Viral loads of these samples were then measured by a RT-qPCR assay resulting in data consistent with those obtained by the diagnosis assay kit (Table 1 and Figure 3). Accordingly, the highest viral loads were obtained from those specimens that allowed us to isolate ZIKV by Vero cell infections. The urine of patient 1 exhibited a ZIKV-genomic RNA copies of 2.68 × 10^3^ per ml whereas the patient 6 displayed 2.53 × 10^s^ ZIKV RNA copies per ml in urine and 7.44 × 10^4^ ZIKV RNA copies per ml in saliva. As expected for isolated viral samples, we observed an increase of genomic ZIKV RNA copies in Vero-cell- isolated samples, in which the isolated from patient 1 presented 1.24 × 10^10^ copies/ml and patient 6, 2.88 × 10 ^9^ copies/ml (Figure 3). Furthermore, we confirmed positivity of the urine from patient 7 and the positive detection of ZIKV RNA in saliva and urine of patient 9, although this established value is borderline localized in the limit of detection.

**Fig 3.**
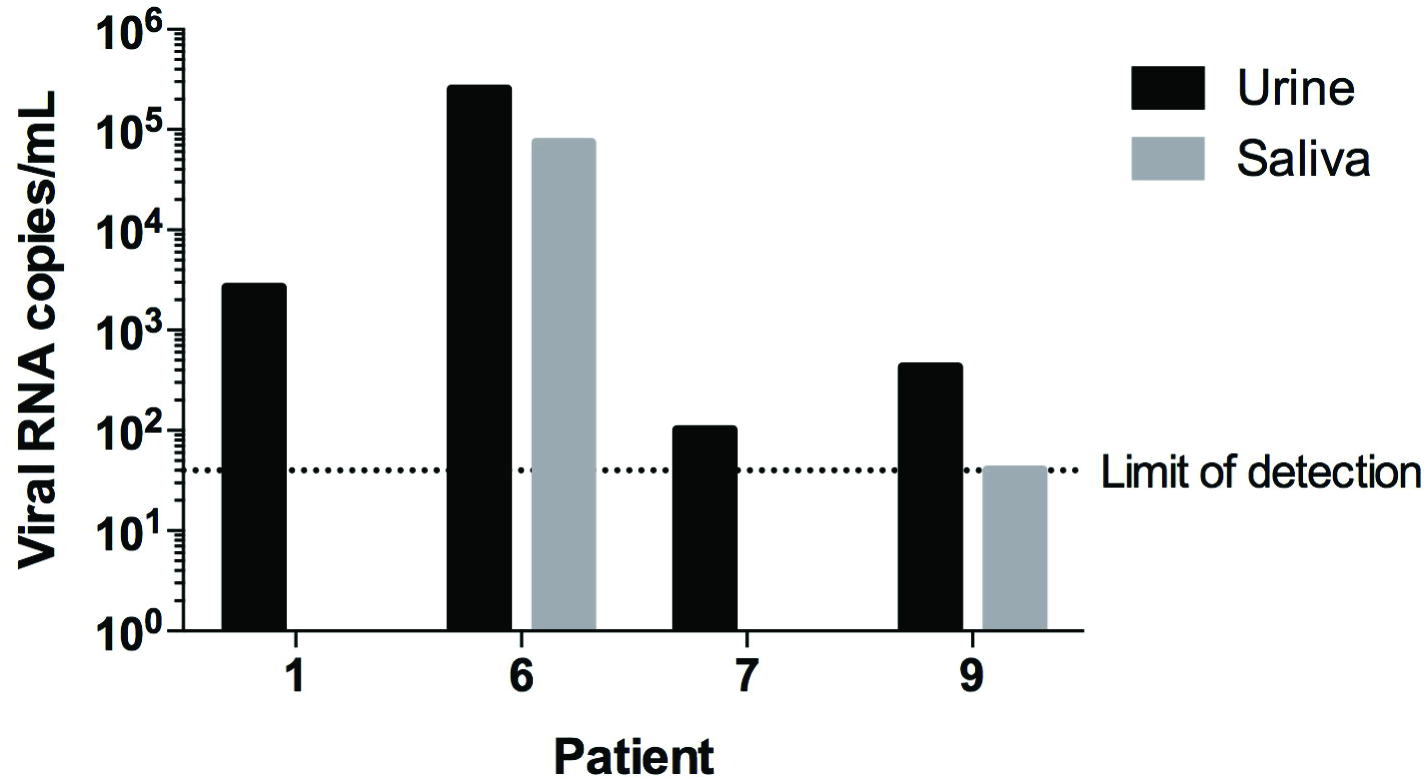
ZIKV Viral loads from urine and saliva specimens of infected patients measured by RT-qPCR.

Urine specimens are shown in black and saliva specimens are shown in grey. The limit of detection is shown as a dotted line corresponding to 40 viral RNA copies/mL.

### ZIKV genomic sequencing

The genomic sequences of Vero cell isolates ZIKV Rio-U1 strain (KU926309), isolated from urine and Rio-S1 (KU926310) strain, isolated from saliva, were then determined. The comparison between Rio-U1 and Rio-S1 yielded 99.61% identity, displaying six amino acid variations in the viral proteins (Table 2). For phylogenetic analysis, we used nucleotide sequences coding the complete ZIKV polyprotein. We observed that all sequences sampled in the Americas form a robust monophyletic cluster (bootstrap score = 97%) within the Asian genotype and share a common ancestor with the ZIKV strain that circulated in French Polynesia in November 2013 and remained genetically isolated from African clusters (Fig 4).

**Fig 4.**
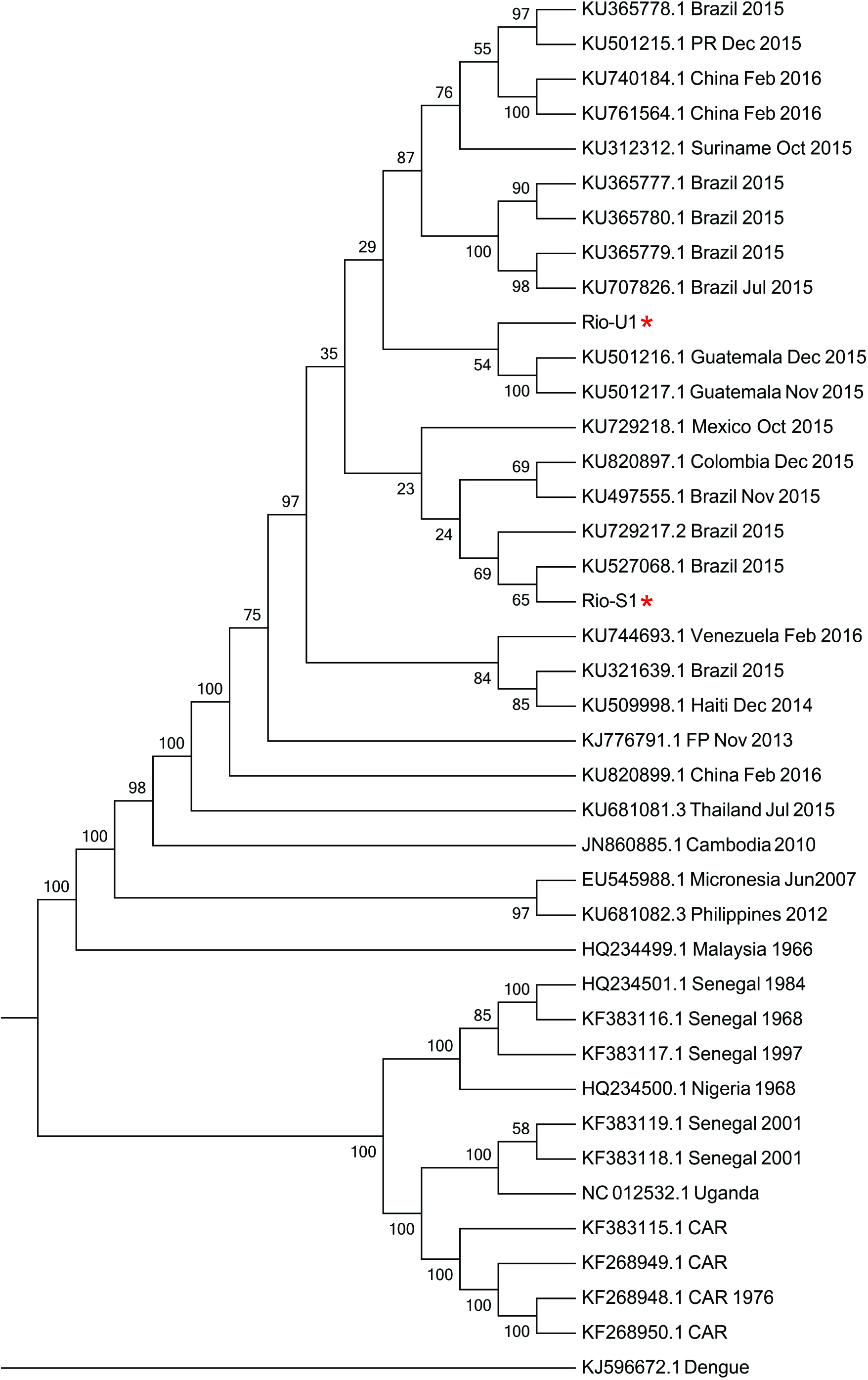
Molecular Phylogenetic analysis by Maximum Likelihood method.

The evolutionary history was inferred by using the Maximum Likelihood method based on the General Time Reversible model. The bootstrap consensus tree inferred from 1000 replicates is taken to represent the evolutionary history of the taxa analyzed. Branches corresponding to partitions reproduced in less than 50% bootstrap replicates are collapsed. The percentage of replicate trees in which the associated taxa clustered together in the bootstrap test (1000 replicates) are shown next to the branches. Initial tree(s) for the heuristic search were obtained automatically by applying Neighbor-Join and BioNJ algorithms to a matrix of pairwise distances estimated using the Maximum Composite Likelihood (MCL) approach, and then selecting the topology with superior log likelihood value. A discrete Gamma distribution was used to model evolutionary rate differences among sites (5 categories (+G, parameter = 0.9645)). The rate variation model allowed for some sites to be evolutionarily invariable ([+I], 37.8665% sites). The analysis involved 40 nucleotide sequences. All positions with less than 95% site coverage were eliminated. That is, fewer than 5% alignment gaps, missing data, and ambiguous bases were allowed at any position. There were a total of 10247 positions in the final dataset. Evolutionary analyses were conducted in MEGA7

Phylogenetic analysis of the isolated viruses exhibiting the highest identity of ZIKV strain Rio-U1 with KU501216.1 and KU501217.1 both from Guatemala (99.7 % identity), isolated also related with the first reported autochthonous transmission of ZIKV in Brazil [31]. Whereas Rio-S1 presented 99.7% of identity with KU527068.1, isolated in Brazil from a Zika-associated microcephaly case [14].

## DISCUSSION

In this study, we demonstrate the occurrence of infectious Zika viral particles in urine and saliva of patients. Besides, we also showed that the saliva of an acute phase patient may have a viral concentration of 80 PFU/ml. The isolation of two ZIKV samples from urine and saliva was associated with ZIKV load in infected patients during the acute phase. Actually, the presence of ZIKV genome in urine is not a novelty. Hence, former studies preconized the use of urine and saliva for ZIKV RNA detection and diagnosis [27, 30], since ZIKV genome was more frequently identified in saliva and urine compared to blood. Furthermore, the finding of flaviviral genome in urine was earlier described in Dengue [32], Yellow Fever [33], St. Louis Encephalitis [34], Japanese Encephalitis [35], and West Nile viruses [36]. Dengue genome was also detected in saliva of infected patients [32]. Interestingly, the existence of excreted-infectious West Nile particles in the urine of acute phase patients was earlier described in conjunction with their isolation in Vero E6 and in BHK21 cells [37]. Particularly, ZIKV isolation was approached by many groups utilizing Vero cells (GeneBank: KJ776791; JN860885; KU647676). Therefore, we adopted this cell model to detect, amplify and quantify viable ZIKV straight from patient’s samples of urine and saliva.

The recovery of ZIKV from these urine and saliva was effective in two of nine patients whose viral load were clearly detectable. Interestingly, despite the fact that the viral load found in the urine of patient 1 was considerably lower, around one hundred times, than the equivalent sample in patient 6, we only recovered virus from urine of the former (Rio-U1 strain). On the other hand, recovering of infective ZIKV from patient 6, the Rio-S1 strain, was successful using the saliva sample, but not with urine one, even though the highest number of copies has been established in urine. Concordantly, we detected in this analysis a superior number of plaques in plaque assay of saliva. Viral detection and recovery from urine and saliva of ZIKV patients might be firstly related to the severity of infection as well as the period of specimen collection after the onset of Zika symptoms. The detection of ZIKV RNA in saliva improved the diagnosis in the first week from the disease onset [30]. But ZIKV viruria persists for longer periods after disease beginning and, in some cases, for longer than two weeks from Zika onset [27], as described in the two recently reported cases of Guillain–Barré syndrome occurred in Martinica [38]. However, it is necessary to perform additional clinical studies associating disease onset, severity of symptoms and viral persistence in urine and saliva to better clarify this point.

Another aspect in viral recovering deals with the physiological pH found in saliva and urine. Hence, pH in urine varies from 4.5 to 8.0 while saliva assumes values near neutral pH. It is well known that the flavivirus envelope protein E undergoes irreversible conformational changes at a mildly acidic pH (below 6.5), a process naturally occurring in the viral membrane fusion in endosomes [39]. These structural changes are irreversible, and outside of cellular environment, provoke loss of infectivity and hemagglutination activity as well as virus aggregation due to increased hydrophobicity [40]. Thereby, we suggest that the failure of recovering ZIKV strain in Vero cells propagation from the urine of patient 6 would be due to the inactivation of most ZIKV due to exposition of the acidic pH value of 5.6 of this urine specimen. The infectious virus number was lower, at least proportionally to the viral RNA copies presented in this fluid, when compared to saliva of the same patient. We do not establish the pH of patient’s 1 urine, due to volume sample limitations. The importance of ZIKV in urine for human transmission is unexplored, but the effect of acidic pH on viral viability might represent a serious restriction for viral spreading. In West Nile Virus when a similar urine excretion occurs, it is considered that the presence of infectious particles would represent a real risk for inter human transmission through kidney transplantation [37].

In reference to the occurrence of viable ZIKV in saliva, a large range of viruses can be identified in this specimen, such as Cytomegalovirus, Ebola virus, Enteroviruses, Hepatitis B virus, Hepatitis C virus, Human herpesviruses, HIV, Human papillomavirus, Influenza virus, Measles virus, Rhinoviruses and Rubella virus [41, 42]. As previously mentioned, Zika and dengue virus were also discovered in saliva [30, 32]. Although, the presence of intact viral particles in saliva do not distinguish viable virus from noninfectious virus. However, for the first time, we could well identify ZIKV plaque forming units from saliva of an infected man in Vero cell monolayers with a titer corresponding to 80 PFU per ml.

Essentially, another important subject is that the existence of viable virus in oral fluid samples does not always indicate that the virus can be transmitted orally and become epidemiologically relevant. Actually, viral infections of the oral cavity are relatively rare, since saliva contains antiviral molecules and is relatively hypotonic being capable of lysing enveloped viruses [43]. Perhaps, the established proportion of approximately 1 PFU to 1,000 ZIKV RNA copies in saliva of one patient was modulated by these host factors.

Although saliva functions as a protective barrier for virus entry, some studies have shown that a disruption in oral mucosa or periodontal disease can facilitate virus entry [44]. Since previous studies detected Flaviviruses as Dengue [45, 46] and Zika [30] virus in saliva, and our study have demonstrated possible infectious ability of Zika viral particles in saliva, a potential person-to-person Zika virus infection through this specimen, using a disrupted oral mucosa or periodontal pockets as virus entry, should be considered and investigated.

ZIKV is an emergent vector-borne disease, but fast growing evidence points to an increased relevance of its non-vector ways of transmission, as perinatal and transplacental transmission occurs from mother to child [14, 23]. Additionally, ZIKV genome was also detected in breast milk, followed by viral isolation of infective viral particles [29]. Moreover, cases of probable sexual transmission have been reported with association of ZIKV in semen [25, 26]. In addition, viral contamination linked to blood transfusion and organ transplantation have been previously discussed [47]. Furthermore, reports of laboratorial infection or bites of animals was associated to the transmission [48]. Finally, evidence of vertical and/or venereal transmission between mosquitoes was supported by the detection of ZIKV natural infection in males *Ae. furcifer* [19].

We compared the complete coding sequences obtained in this study with public sequence data from Zika virus representative of the isolates from three distinct genotypes in Asian, West African, and East African in addition to isolates from recent outbreak in Americans. Similarly to the sequences described in the recent widespread epidemic of ZIKV in the Americas, the sequences Rio-S1 and Rio-U1 from ZIKV isolated in this study clustered with the Asian clade, covering sequences from New World, Pacific, Micronesian and Malaysian strains.

Since surveillance programs have reported periodic circulation of the ZIKV virus since 1968, with high frequency activity varying an interval of 1 – 2 years added to fact that RNA virus evolve fast, their host and vector broad range, non-vector transmission, and particularly risk of neurotropic and teratogenic outcomes, the molecular epidemiologic vigilance is crucial to solve this questions.

In conclusion, the detection of infective ZIKV in saliva and urine of patients deserves a more detailed study to establish whether or not these fluids contribute to viral transmission. Surely, these findings will be extremely relevant to prevent and control ZIKV transmission.

## Acknowledgments

The authors thank Denise Valle from Laboratório de Biologia Molecular de Flavivírus ‐IOC/FIOCRUZ - RJ for for helpful discussion and comments about the manuscript and supporting our studies on vector-borne-diseases. To Lais de Souza Soares from Laboratório de Biologia Molecular de Flavivírus - IOC/FIOCRUZ - RJ, Anielly Alves Ferreira and Rafaela Miranda from Laboratório de Mosquitos Transmissores de Hematozoários - IOC/FIOCRUZ - RJ, Aline dos Santos Moreira, Renata Almeida de Sá and Beatriz de Lima Alessio Müller from Plataforma Genômica-Sequenciamento de DNA/RPT01A/FIOCRUZ for technical support and Ricardo Magrani Junqueira from Plataforma de Sequenciamento de Alto Desempenho/IOC-FIOCRUZ – RJ, for sharing expertise and information on NGS. We are grateful to Miriã Alves Gonçalves Trindade, Rosiléa de Souza Rosa dos Santos, Mirian da Conceição Sobrinho, Tulanne Yuriê da Silva Santos from Instituto Nacional de Infectologia - Ambulatório de Doenças Febris Agudas - INI/FIOCRUZ – RJ for helping with samples and data collection, Heloísa Diniz from Serviço de Multimeios/CICT-FIOCRUZ and Gutemberg Brito from SEJOR/IOC-FIOCRUZ for the design of photographs and figures. We also thank Ana Paula de Campos Guimarães from Unidade Genömica Computacional Darcy Fontoura/ LNCC and Guilherme Loss and Rafael Guedes from Laboratòrio Nacional de Bioinformática/ LNCC for *De novo* sequencing, assembly and analysis of the genome of the Zika Virus.

